# Personalized single-cell networks: a framework to predict the response of any gene to any drug for any patient

**DOI:** 10.1101/837807

**Authors:** Haripriya Harikumar, Thomas P. Quinn, Santu Rana, Sunil Gupta, Svetha Venkatesh

## Abstract

**Background:** The last decade has seen a major increase in the availability of genomic data. This includes expert-curated databases that describe the biological activity of genes, as well as high-throughput assays that measure gene expression in bulk tissue and single cells. Integrating these heterogeneous data sources can generate new hypotheses about biological systems. Our primary objective is to combine population-level drug-response data with patient-level single-cell expression data to predict how any gene will respond to any drug for any patient.

**Methods:** We take 2 approaches to benchmarking a “dual-channel” random walk with restart (RWR) for data integration. First, we evaluate how well RWR can predict known gene functions from single-cell gene co-expression networks. Second, we evaluate how well RWR can predict known drug responses from individual cell networks. We then present two exploratory applications. In the first application, we combine the Gene Ontology database with glioblastoma single cells from 5 individual patients to identify genes whose functions differ between cancers. In the second application, we combine the LINCS drug-response database with the same glioblastoma data to identify genes that may exhibit patient-specific drug responses.

**Conclusions:** Our manuscript introduces two innovations to the integration of heterogeneous biological data. First, we use a “dual-channel” method to predict up-regulation and down-regulation separately. Second, we use individualized single-cell gene co-expression networks to make personalized predictions. These innovations let us predict gene function and drug response for individual patients. Taken together, our work shows promise that single-cell co-expression data could be combined in heterogeneous information networks to facilitate precision medicine.

## 1 Introduction

Advances in high-throughput RNA-sequencing (RNA-Seq) have made it possible to quantify RNA presence in any biological sample [29], producing a gene expression signature that can serve as a biomarker for disease prediction [15, 1, 47] or surveillance [31, 38]. Over the last few years, single-cell RNA-Seq has risen in popularity [14]. Compared with conventional bulk RNA-Seq, which measures the average gene expression for an individual sample, single-cell RNA-Seq (scRNA-Seq) measures gene expression for an individual cell. This new mode of data makes it possible to explore tissue heterogeneity, notably tumor heterogeneity [25], by producing multiple data points per individual (i.e., one for each cell). Since genes are often understood to work in cooperative modules, the analysis of *gene co-expression networks* is commonplace. For bulk RNA-Seq, a gene co-expression network describes how genes co-occur for a population of individual samples. For scRNA-Seq, the network describes gene co-expression for a population of single cells. When these cells belong to an individual patient, the scRNA-Seq network is a kind of **personalized network** that one could use for precision medicine tasks.

Gene co-expression networks can be integrated with outside information to combine **general knowledge** (in the form of a relational database like Gene Ontology [2]) with **specific knowledge** about a sample (in the form of a co-expression network). For example, weighted gene co-expression network analysis is a popular method for functionally characterizing parts of the network, or the network as a whole [23, 24]. Although these coarse descriptions are useful, one could also combine general- and specific knowledge to make finer-level predictions about the behavior of *individual genes*. By representing each modality as a graph, multiple data streams can be combined into a **heterogeneous information network**, and then analyzed under a unified framework based on the principle of “guilt-by-association” [45] (e.g., if “a” is connected to “b” and “b” is connected to “c”, then “a” is probably connected to “c). When the general knowledge is **gene-annotation** associations, we can (a) impute the function for genes with no known role or (b) select the most important known function. When the general knowledge is **gene-drug** response, we can predict the response of any gene to any drug. Since these inferences are tailored to the co-expression network used, they can be made personalized by using the single-cell network of an individual patient.

Random walk (RW) is a popular method that offers a general solution to the analysis of heterogeneous information networks [34, 45]. There are many variants to RW, including random walk with restart (RWR), where each step has a probability of restarting from the starting node (or a neighbor of the starting node) [44]. RW and RWR are often used in recommendation systems [3, 8, 19], but can also perform other machine learning tasks like image segmentation [16, 18], image captioning [32], or community detection [36, 22]. One advantage of RW is that it can handle missing data [17], making it a good choice for processing sparse gene annotation databases. RW and RWR have both found use in biology to find associations between genes and another data modality. For example, the “InfAcrOnt” method used an RW-based method to infer similarities between ontology terms by integrating annotations with a gene-gene interaction network [6]. Similarly, the “RWLPAP” method used RW to find lncRNA-protein associations [50], while others have used RW to predict gene-disease associations [51]. Meanwhile, RWR has been used to identify epigenetic factors within the genome [26], key genes involved in colorectal cancer [9], novel microRNA-disease associations [43], infection-related genes [52], disease-related genes [46], and functional similarities between genes [35]. Bi-random walk, another random walk variant, has been used to rank disease genes from a protein-protein interaction network [48].

In contrast to the previous works, which make use of population-level graphs, *we apply RWR to patient-level graphs, allowing us to make predictions about gene behavior that are personalized to each patient*. We take 2 approaches to benchmarking “dual-channel” RWR for data integration. First, we evaluate how well RWR can predict known gene functions from single-cell gene co-expression networks. Second, we evaluate how well RWR can predict known drug responses from individual cell networks. We then present two exploratory applications. In the first application, we combine the Gene Ontology database with glioblastoma single cells from 5 individual patients to identify genes whose functions differ between cancers. In the second application, we combine the LINCS drug-response database with the same glioblastoma data to identify genes that may exhibit patient-specific drug responses. Taken together, our work shows promise that single-cell co-expression data could be combined in heterogeneous information networks to facilitate precision medicine.

## 2 Methods

### 2.1 Overview

In the medical domain, gene expression can be used as a biomarker to measure the functional state of a cell. One way in which drugs mediate their therapeutic or toxic effects is by altering gene expression. However, the assays needed to test how gene expression changes in response to a drug can be expensive and time consuming. Imputation has the potential to accelerate research by “recommending” novel gene-drug responses. Random walk methods can combine sparse heterogeneous graphs based on the principle of “guilt-by-association” [45]. Figure 1 provides an abstracted schematic of the proposed framework. Figure 2 provides a visualization of the input and output for the random walk with restart (RWR) method. Figure 3 presents a bird’s-eye view of the data processing, validation, and application steps performed in this study.

**Figure 1:**
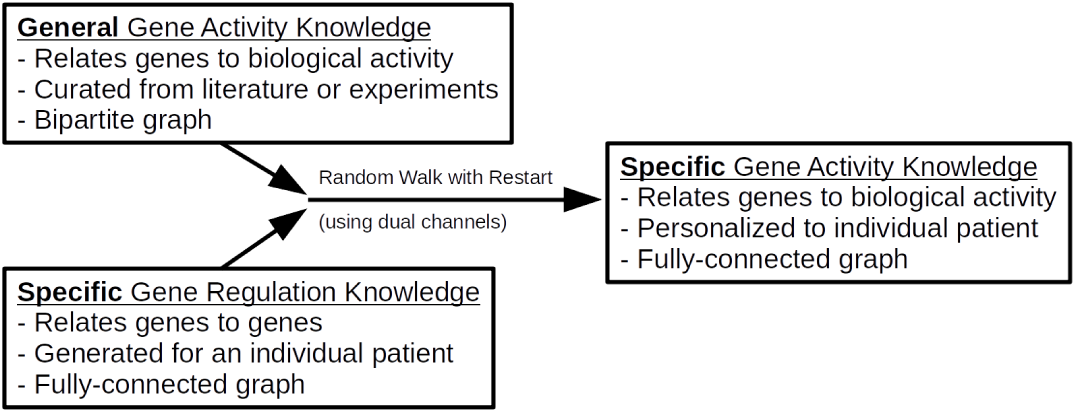
An abstracted schematic of the proposed framework. Expert-curated databases like Gene Ontology (GO) and the Library of Integrated Network-Based Cellular Signatures (LINCS) can provide some **general knowledge** about biological activity. High-throughput single-cell sequencing assays can provide **specific knowledge** for an individual patient. Random walk with restart (RWR) can combine these heterogeneous data sources to provide specific knowledge about biological activity for an individual patient. This framework allows us to predict how any gene will respond to any drug for any patient.

**Figure 2:**
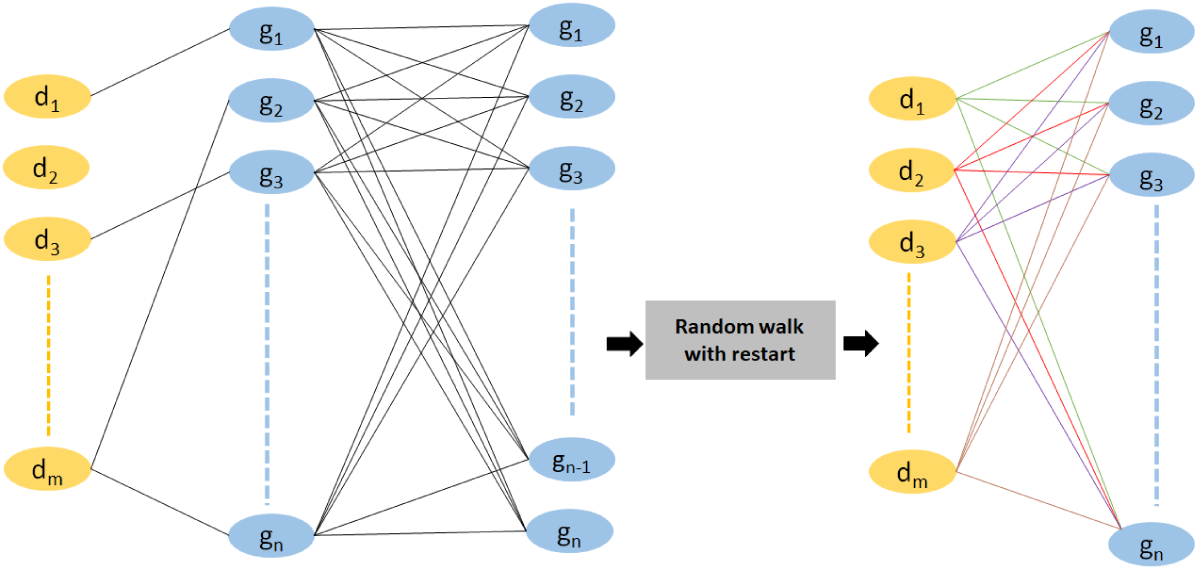
Our goal is to combine a generic gene-based bipartite graph with the auxiliary knowledge of a fully-connected personalized graph. RWR will impute the missing links (and update the existing links) by “walking through” the auxiliary information. The left panel shows a (sparsely-connected) gene-drug graph combined with a (fully-connected) gene-gene graph, where *d*_*i*_ represents the drugs and *g*_*i*_ represents the genes. The example gene-drug network has missing links. The right panel shows the output of RWR: a complete network of newly predicted gene-drug interactions. Here, the missing link between any drug *d*_*i*_ and any gene *g*_*i*_ is replaced with a new link. The method works based on the principle of “guilt-by-association”: the value of the new *d*_*i*_−*g*_*i*_ links will be large if *g*_*i*_ is strongly connected to genes that are also connected to *d*_*i*_. When the gene-drug graph is fully-connected, RWR will instead “update” the importance of each connection.

**Figure 3:**
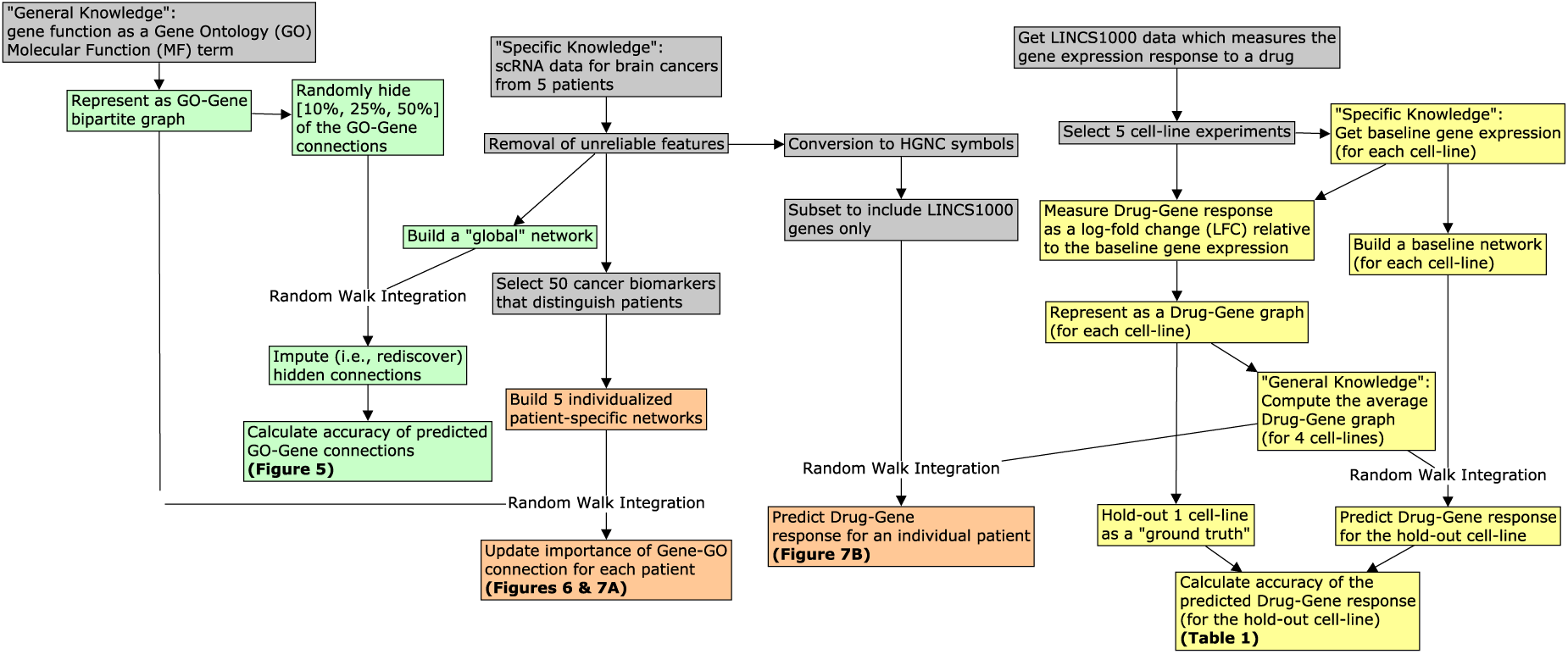
A bird’s-eye view of the data collection, integration, and analysis steps performed in this study. We use RWR to combine general knowledge with some specific knowledge about a sample. We separately use gene function and drug-response data as the source of general knowledge. We use co-expression networks as the source of specific knowledge. By combining the drug-response data with an individualized single-cell network, we can make predictions about gene behavior that are personalized to each patient. Concept nodes are colored by activity type: data processing (gray), validation of gene-annotation prediction (green), validation of gene-drug prediction (yellow), and exploratory application (orange).

### 2.2 Data acquisition

The gene expression data come from two primary sources. First, we acquired single-cell RNA-Seq (scRNA-Seq) expression data for 5 glioblastoma multiforme tumors [33] using the recount2 package for the R programming language [7] (ID: SRP042161). Since scRNA-Seq data are incredibly sparse, and since the random walk with restart algorithm is computationally expensive, we elected to remove genes that had zero values in more than 25% of cells. This resulted in 3022 genes. Finally, we randomly split the cells into 5-folds per patient so that we could estimate the variability of our downstream analyses. Second, we acquired gene expression data from the Library of Integrated Network-Based Cellular Signatures (LINCS) [20] using the Gene Expression Omnibus (GEO) [10] (ID: GSE70138). We split these LINCS data into smaller data sets based on the cell line ID under study. We *a priori* included the A375 (skin; malignant melanoma), HA1E (kidney; embryonic), HT29 (colon; adenocarcinoma), MCF7 (breast; adenocarcinoma), and PC3 (prostate; adenocarcinoma) cell lines because they were treated with the largest number of drugs.

### 2.3 Defining the gene co-expression network graphs

Although correlation is a popular choice for measuring gene co-expression, correlations can yield spurious results for next-generation sequencing data [27]. Instead, we calculate the proportionality between genes using the *ϕ*_*s*_ metric from the propr package for the R programming language [39]. Although this does not offer a perfect solution [12], studying gene-gene proportionality has a strong theoretical justification [27] and empirically outperforms other metrics of association for scRNA-Seq [41].

The proportionality metric describes the dissimilarity between any two genes, and ranges from [0, ∞), where 0 indicates a perfect association. We converted this to a similarity measure *ϕ*_*i*_ that ranges from [0, 1] by max-scaling *ϕ*_*i*_ = (max(*ϕ*_*s*_)*‐ϕ*_*s*_)*/*max(*ϕ*_*s*_), such that *ϕ*_*i*_ = 1 when *ϕ*_*s*_ = 0. A gene-gene matrix of *ϕ*_*i*_ scores is analogous to a gene-gene matrix of correlation coefficients, and constitutes our gene co-expression network. We calculated the *ϕ*_*i*_ co-expression network for the entire scRNA-Seq data set (1 network), for each of the 5-folds per-patient (25 networks total), and for each LINCS baseline (drug-free) cell line (5 networks total). All co-expression networks are available from https://zenodo.org/record/3522494.

### 2.4 Defining the bipartite graphs

We constructed two types of bipartite graphs: the **gene-annotation graph** and the **gene-drug graph**. First, we made the gene-annotation graph from the Gene Ontology Biological Process database [2] via the AnnotationDbi and org.Hs.eg.db Bioconductor packages. An edge exists whenever a gene is associated with an annotation. Second, we made the gene-drug graphs using the LINCS data. For each cell line, we computed a gene-drug graph by calculating the log-fold change between the median of the drug-treated cell’s expression and the median of the drug-naive cell’s expression. This results in a fully-connected and weighted bipartite graph, where a large positive value means that the drug causes the gene to up-regulate (and *vice versa*). All bipartite graphs are available from https://zenodo.org/record/3522494.

### 2.5 The combined co-expression and bipartite graph

Consider a graph *G* with *V* = 1…*N* vertices, *E*^+^ positive edges, and *E*^−^ negative edges. The graphs used for our analyses are composed for two parts: a (general knowledge) bipartite graph and a (specific knowledge) fully-connected gene co-expression graph. For a bipartite graph, the vertex set *V* can be separated into two distinct sets, *V*_1_ and *V*_2_, such that no edges exist within either set. For a fully-connected (or complete) graph, there exists an edge between every pair of vertices within one set. For the graph *G*, the bipartite and fully-connected graphs are joined via the common vertex set *V*_1_ that contains genes and *V*_2_ contains annotations or drugs.

### 2.6 Dual-channel random walk with restart (RWR)

Traditional RWR methods can only perform a random walk on graphs with positive edge weights [34]. Since the response of a gene to a drug is directional (up-regulated or down-regulated), we chose to use a modified RWR method, proposed by [5], that handles graphs with both positive and negative edge weights. Random walk requires transition probability matrices to decide the next step in the walk. The Chen et al. transition probability matrices can be computed based on the following equations:

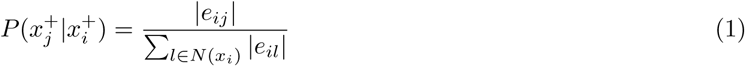

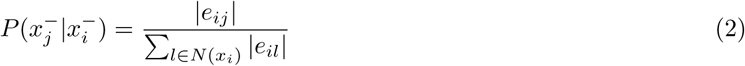

when *e*_*ij*_ ≥ 0, and

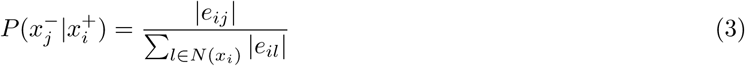

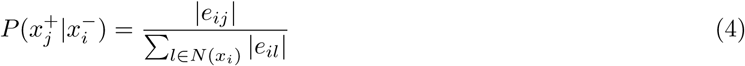

when *e*_*ij*_ < 0. For all equations, *e*_*ij*_ is the edge weight between nodes *x*_*i*_ and *x*_*j*_, and *N* (*x*_*i*_) is the set of neighbors for node *x*_*i*_. These equations separate out the positive (and negative) transitions, and are used to calculate the total positive (and negative) information flow for each node. They are fixed for all steps.

Though the transition probabilities are computed separately, the information accumulated in a node depends on both the positive and negative information which flows through the node. For example, the positive information in a node depends on the negative information of any neighboring node connected by a negative edge weight (think: negative times negative is positive). Likewise, negative information in a node depends on the positive information in a neighboring node connected by a negative edge weight, and *vice versa* (think: negative times positive is negative). Figure 4 illustrates the information flow to a node *x*_*j*_ from two neighbors. The flow of information between the positive “plane” of the graph to the negative “plane” of the graph can be formulated with the equations:

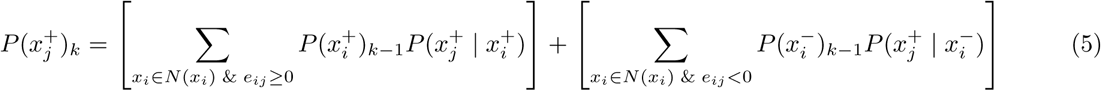

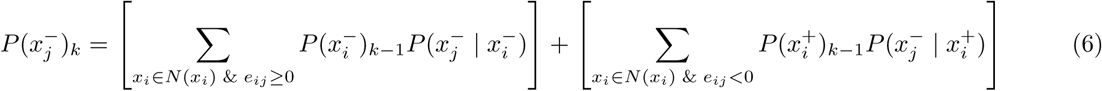

where the probability 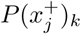 is updated at each step *k* = 2…10000.

**Figure 4:**
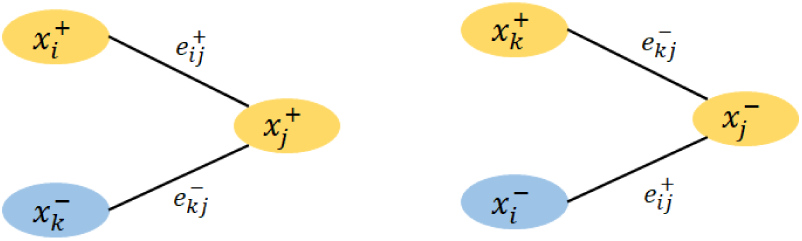
This figure illustrates the flow of information between adjacent nodes. The positive information nodes are yellow and the negative information nodes are purple. A positive edge weight is represented by 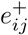, while a negative edge weight is represented by 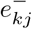. The sign of the edge weights determines which information (positive or negative) flows from one node to another. The positive information of a node *x*_*j*_ depends on the positive information of *x*_*i*_ when the edge is positively weighted (think: positive times positive is positive). The negative information of a node *x*_*j*_ depends on the negative information of *x*_*i*_ when the edge is positively weighted (think: negative times positive is negative). Similarly, the negative flow of negative information can contribute to positive information. The “dual-channel” RWR algorithm incorporates these edge weights. Note that in practice each node will contain both positive and negative information.

RWR always considers a probability *α* to return back to the original nearest neighboring nodes at each step in the random walk. This is used to weigh the importance of node-specific information with respect to the whole graph, including for long walks:

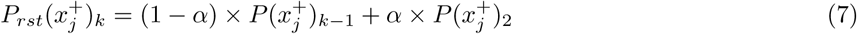

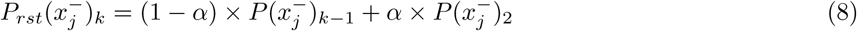

where the restart probability 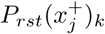 is updated at each step *k* = 2…10000, and 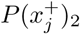 is the probability after the first update. These equations find the positive and negative restart information with respect to the node *x*_*j*_. Each 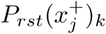 is a vector of probabilities that together sum to 1. This probability has two parts: the global information and the local information. The local information is the initial probability with respect to the nearest neighbors of node *x*_*j*_, and is denoted by 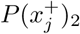 [or 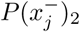] (i.e., the probability after the first update). The restart probability *α* is chosen from the range [0, 1], where a higher value weighs the local information more than the global information. We chose *α* = 0.1 to place a larger emphasis on the global information. Simulations with a toy data set verified this choice.

### 2.7 Analysis of random walk with restart (RWR) scores

For each gene, the RWR algorithm returns a vector of probabilities that together sum to 1. According to the guilt-by-association assumption, we interpret these probabilities to indicate the strength of the connection between the reference gene and each target. Since we are only interested in gene-annotation and gene-drug relationships, we exclude all gene-gene probabilities. Viewing the probability distribution as a composition (c.f., [11]), we perform a centered log-ratio transformation of each probability vector subset. This transformation normalizes the probability distributions so that we can compare them between samples [4, 37]. We define the RWR score 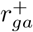 (or 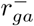) for each gene-annotation connection as the transform of its RWR probability:

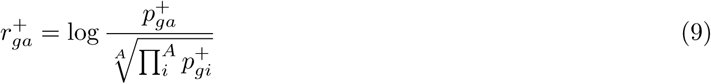

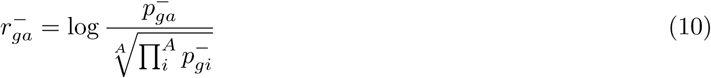

for a bipartite graph describing *g* = 1…*G* genes and *a* = 1…*A* annotations (or *A* drugs), where 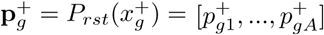 (i.e., from the final step). These transformed RWR scores can be used for univariate statistical analyses, such as an analysis of variance (ANOVA) (c.f., analysis of compositional data [13, 28]).

### 2.8 Benchmark validation

#### 2.8.1 Validation of gene-annotation prediction

Our strategy to validate RWR for gene-annotation prediction involves “hiding” known functional associations and seeing whether the RWR algorithm can re-discover them. This is done by turning 1s into 0s in the bipartite graph, a process we call “sparsification”. Our sparsification procedure works in 4 steps. First, we combine the original GO BP (or MF) bipartite graph with the master single-cell co-expression graph. Second, we subset the graph to include 25% of the gene annoations and 25% of the genes (this is done to reduce the computational overhead). Third, we randomly hide [10, 25, 50] percent of the gene-annotation connections from the bipartite sub-graph. Since this random selection could cause a feature to lose all connections, we use a constrained sampling strategy: the subsampled graph must contain at least one non-zero entry for each feature. Fourth, we apply the RWR algorithm to the sparsified and non-sparsified graphs, separately. We repeat this process 25 times, using a different random graph each time. By comparing the RWR scores between the hidden and unknown connections, we can determine whether our method rediscovers hidden connections.

#### 2.8.2 Validation of gene-drug prediction

We use a different strategy to validate RWR for drug-response prediction. Since we have the gene-drug and gene-gene interaction data for 5 cell lines (A375, HA1E, HT29, MCF7 and PC3), we can set aside the known gene-drug responses for 1 cell line (PC3) as a “ground truth” test set. Then, we can use a composite of the remaining 4 gene-drug graphs to predict the gene-drug responses for the withheld cell line.

This is done in two steps. First, we use the averaged gene-drug data for 4 cell lines (a general drug graph) and the gene-gene data for PC3 (a specific gene graph) to impute the gene-drug response for PC3 (a specific drug graph). In the second step, we use the gene-drug data for PC3 (a specific drug graph) and its corresponding gene-gene data (a specific gene graph) to calculate the “ground truth” RWR scores for PC (a specific drug graph). The “ground truth” is the RWR scores when all PC3 drug-response experiments have been performed. With these two outputs, we can calculate the agreement between the imputed and “ground truth” RWR scores (using Spearman’s correlation, MSE, and accuracy).

### 2.9 Exploratory application of gene-drug prediction

Having demonstrated that RWR can perform well for single-cell co-expression networks, and can make meaningful drug-response predictions from composite LINCS data, we combine these heterogeneous data sources to make personalized drug-response predictions for individual single-cell networks. This requires some data munging. First, we transform the ENGS features used by the single-cell data into the HGNC features used by LINCS (only including genes with a 1-to-1 mapping, resulting in 181 genes). Second, we build an HGNC co-expression network with *ϕ*_*i*_ (for 5 folds of 5 patients, yielding 25 networks total). Third, we combine the composite LINCS gene-drug bipartite graph with each of the 25 HGNC single-cell networks. Fourth, we use our RWR algorithm to predict how 181 genes would respond to 1732 drugs for each patient fold. As above, we perform an analysis of variance (ANOVA) to detect inter-patient differences.

## 3 Results and Discussion

### 3.1 Why use single cells?

In this study, we analyze a previously published single-cell data set that measured the gene expression for 5 glioblastoma patients. A principal components analysis of these data show that the major axes of variance tend to group the cells according to the patient-of-origin. Indeed, an ANOVA of gene expression with respect to patient ID reveals that 2204 of the 3022 genes have significantly different expression in at least one patient (FDR-adjusted *p <* .05). This suggests that single-cell gene expression is unique to each patient. Although it is possible to obtain sample-specific *gene expression* using bulk RNA-Seq, our approach to data integration exploits the graphical structure of sample-specific *gene co-expression networks*. scRNA-Seq, generating multiple measurements per individual, makes the computation of sample-specific co-expression networks straightforward (though others have proposed ways to estimate these from bulk RNA-Seq [21, 30]).

### 3.2 Validation of gene-annotation prediction

The Gene Ontology (GO) project has curated a database which relates genes to biological processes (BP) and molecular functions (MF) (called **annotations**). The GO database has widespread use in bioinformatics for assigning “functional” relevance to sets of gene biomarkers [42]. Although GO organizes the semantic relationships between annotations as a directed acyclic graph, we could more simply represent the relationships as a bipartite graph. By combining a (fully-connected) gene co-expression graph with a (sparsely-connected) gene-annotation bipartite graph, RWR can predict new gene-annotation connections.

To test whether the RWR predictions are meaningful, we “hid” a percentage of known gene-annotation links (by turning 1s into 0s in the bipartite graph), and compared the RWR scores for the *hidden* gene-annotation links against those for the *unknown* links (see Methods for a definition of the RWR score). Figure 5 shows that the RWR scores for hidden connections are appreciably larger than for the unknown connections, confirming that RWR can discover real gene-annotation relationships from a single-cell gene co-expression network.

**Figure 5:**
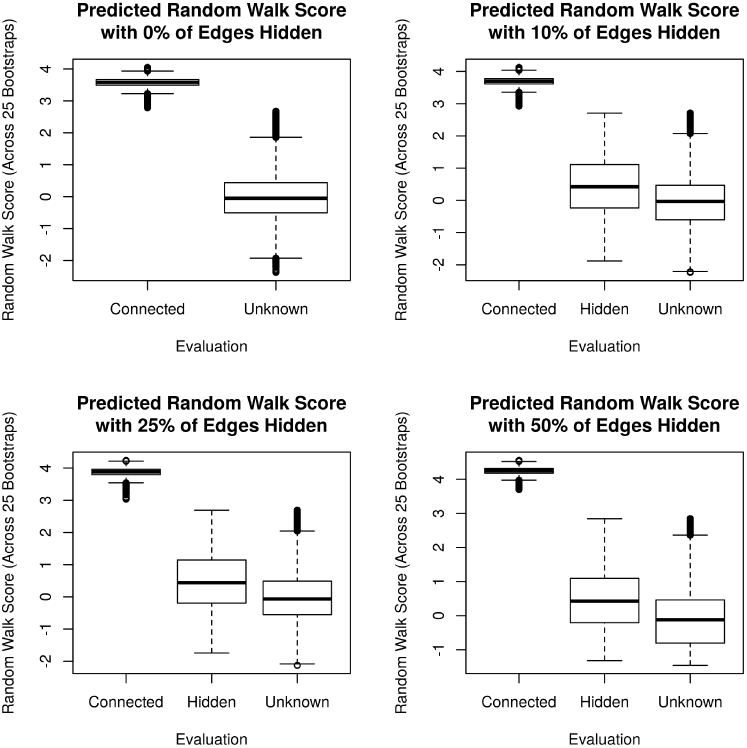
The RWR scores for the hidden and unknown gene-annotation connections (faceted by the amount of sparsity). When known connections are hidden, RWR tends to give higher scores than when connections are unknown, suggesting that RWR can discover real gene-annotation relationships. Note that GO is an incomplete database: the absence of a gene-annotation connection is not the evidence of absence. For this reason, we do know whether the high scoring “Unknown” connections are false positives or previously undiscovered connections. All t-tests comparing “Hidden” and “Unknown” connections have *p* < 10^−15^.

### 3.3 Exploratory application of gene-annotation prediction

Since single-cell RNA-Seq assays measure RNA for multiple cells per patient, we can use these data to build a personalized graph that describes the gene-gene relationships for an individual patient. In order to estimate the variation in these personalized graphs, we divided the cells from each sample into 5 folds (giving us 5 networks per-patient). Above, we show that RWR can discover real gene-annotation relationships. By combining the personalized graph (a kind of *specific knowledge*) with a gene-annotation bipartite graph (a kind of *general knowledge*), the RWR algorithm will score the gene-annotation connections for a given patient. From this, we can identify genes that may have a different functional importance in one cancer versus the others. Taking a subset of the 50 genes with the largest inter-patient differences, we use RWR to compute personalized RWR scores. This results in 25 matrices (for 5 folds of 5 patients), each with 50 rows (for genes) and 369 columns (for BP annotations). Performing an ANOVA on each gene-annotation connection results in a matrix of 50×369 p-values. Figure 6 shows a heatmap of the significant gene-annotation connections (dark red indicates a gene-wise FDR-adjusted *p <* .05). Figure 7A plots the per-patient RWR scores for 4 annotations of the BCL-6 gene that significantly differ between patients. BCL-6 is an important biomarker whose increased expression is associated with worse outcomes in glioblastoma [49]. Our analysis suggests that BCL-6 could have a larger role in inflammation for patients 3 and 5, but a larger role in cartilage development and translational elongation in patient 1. Of course, this hypothesis requires experimental validation.

**Figure 6:**
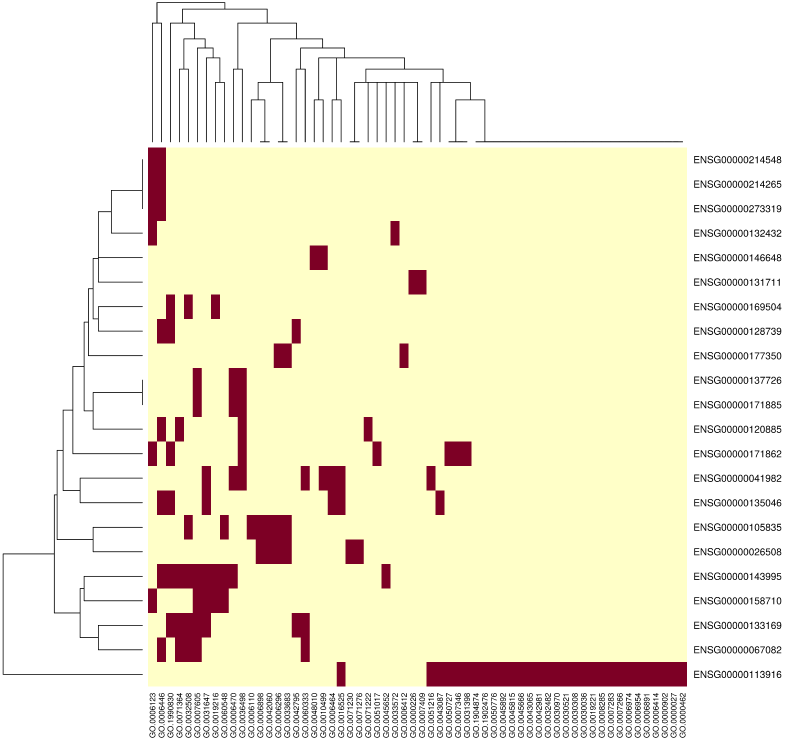
This figure shows a heatmap of the predicted gene-annotation connections that are significantly different between patients (dark red indicates a gene-wise FDR-adjusted *p <* .05). Out of the 50 genes tested, 22 appear to have some form of patient-specific activity.

**Figure 7:**
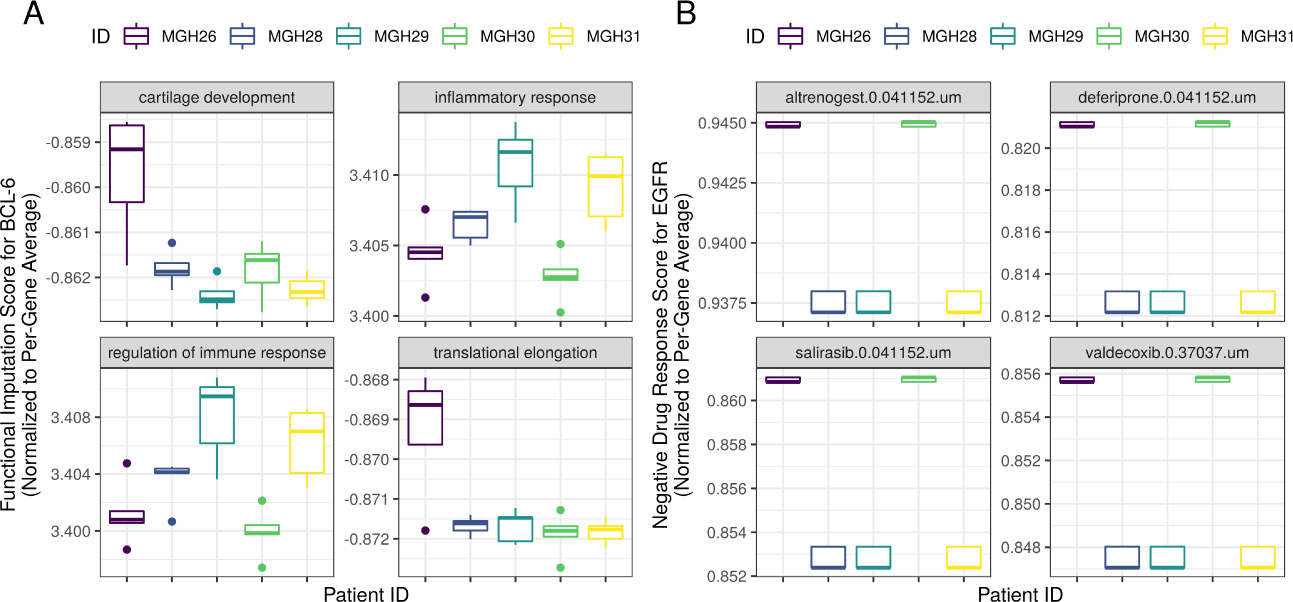
The personalized RWR scores for 4 biological functions of the BCL-6 gene (left panel) and for the EGFR response to 4 drugs (right panel). Panel A suggests that BCL-6 may have a larger role in inflammation for patients 3 and 5, but a larger role in cartilage development and translational elongation in patient 1. Panel B suggests that the anti-inflammatory drug valdecoxib and the anti-neoplastic drug salirasib may cause a stronger down-regulation of EGFR in patients 1 and 4 versus the others.

### 3.4 Validation of gene-drug prediction

The NIH LINCS program has generated a large amount of data on how the gene expression signatures of cell lines change in response to a drug. By conceptualizing the baseline (drug-free) gene co-expression network as a complete graph of *specific knowledge*, and by re-factoring the average gene-drug response as a (weighted) bipartite graph of *general knowledge*, we can apply the same RWR algorithm to predict a cell’s gene expression response to any drug. Since the modified RWR algorithm contains two channels–a positive and negative channel– we can predict up-regulation or down-regulation events separately.

To test whether RWR can make accurate predictions about how a gene in a cell would respond to a drug, we ran the RWR algorithm on the baseline (drug-free) gene co-expression graph of the PC3 cell line using a composite gene-drug graph of 4 different cell lines. We then compared these RWR scores with a “ground truth” (i.e., the RWR scores for when all PC3 drug-response experiments have been performed). The agreement between the composite gene-drug RWR scores and the “ground truth” gene-drug RWR scores tells us how well the composite gene-drug map generalizes to new cell types. Table 1 shows that agreement is high, especially for the top up-regulation and down-regulation events. This confirms that our composite gene-drug graph is useful for drug-response prediction.

**Table 1:**
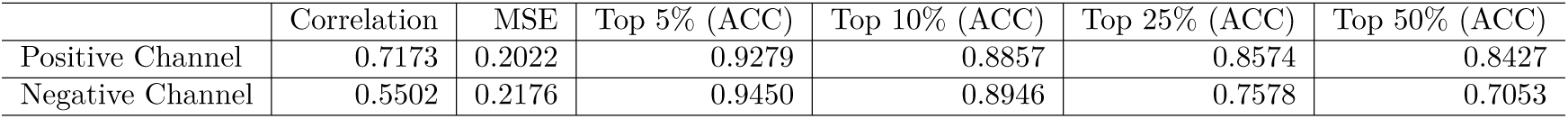
Overall agreement (Spearman’s correlation and MSE) and the accuracy of the overlap (for the top 5%, 10%, 25%, and 50% predicted scores), as calculated separately for the positive and negative channels. Overall, agreement is high, especially for the top up-regulation and down-regulation events.

### 3.5 Exploratory application of gene-drug prediction

The RWR algorithm can combine specific knowledge and general knowledge from disparate sources to make personalized recommendations. This makes RWR a potentially valuable tool for precision medicine.

As an exploratory analysis, we combine the personalized gene co-expression networks with the composite gene-drug graph from LINCS. By running the RWR algorithm on these two data streams, the RWR scores will now suggest how the expression of any gene might change in response to any drug for each of the 5 glioblastoma patients. Using an ANOVA, we identify hundreds of gene-drug connections with RWR scores that differ significantly between patients (gene-wise FDR-adjusted *p <* .05).

Figure 7B shows an example of drugs that have different (negative channel) RWR scores for EGFR. It suggests that the anti-inflammatory drug valdecoxib and the anti-neoplastic drug salirasib may cause a stronger down-regulation of EGFR (a pan-cancer oncogene [40]) in patients 1 and 4 versus the others. The Supplementary Information includes a complete table of the unadjusted ANOVA p-values for the gene-drug inter-patient differences available in https://zenodo.org/record/3743897.

## 4 Limitations

We deployed our framework on only 5 individual patients. As such, we lack a sufficient sample size to test whether any inter-patient differences could be explained by known demographic or clinical phenotypes. It is worth noting that cancer cells are very heterogeneous and, depending on the location of the sample collection, the composition of cell types (and thus gene expression profiles) can change dramatically. As such, factors other than the patient-specific tumour profile could account for differences in the sample-specific gene co-expression networks. Such differences may be difficult to account for without careful experimental design and standardization. Although the scope of this paper is to prove the concept, we wish to remind the readers that much care should be taken when translating the methodology to real-world clinical problem-solving. In the absence of experimental validation, we support our analyses using 2 forms of *in silico* validation, which together demonstrate that RWR can integrate sparse heterogeneous data to discover real biological activity. Although we find the *in silico* validation encouraging, we acknowledge that RWR is merely a prediction tool that recommends hypotheses, and that these predictions may change when the source of general knowledge changes. Experimental validation is needed to determine whether these hypotheses prove true in practice. Further work is needed to validate the clinical relevance of the proposed framework.

## 5 Conclusions

This manuscript describes a framework for combining patient-specific single-cell networks with public drug-response data to make personalized predictions about drug response. Importantly, our approach makes use of a generic framework, and so can be applied to combine many kinds of data. We think the targeted analysis of personalized single-cell networks is promising, and could offer a new direction for precision medicine research.

We conclude with some perspectives on what the future of personalized network analysis may hold. Although RWR can handle sparse heterogeneous data, the positive and negative information obtained for each node can be infinitesimally small. One might address this by first transforming the RWR probabilities. Otherwise, we note that RWR is computationally expensive, making the analysis of high-dimensional data prohibitively slow. One might address this by pre-training a deep neural network to provide an approximate RWR solution. These improvements could help scale personalized predictions to larger graphs.

## Abbreviations

RW: random walk
RWR: random walk with restart
RNA-Seq: RNA sequencing
MSE: mean square error
scRNA-Seq: single-cell RNA sequencing
LINCS: Library of Integrated Network-Based Cellular Signatures
GO: Gene Ontology
BP: Biological Process
MF: Molecular Function
ANOVA: analysis of variance

## Declarations

### Ethics approval and consent to participate

Not applicable.

### Consent for publication

Not applicable.

### Availability of data and material

The raw data are publicly available from the resources described in the Methods. All gene co-expression and bipartite graphs used in these analyses are available from https://zenodo.org/record/3522494.

## Acknowledgements

Not applicable.

## Authors’ contributions

HH implemented the RWR algorithm and applied it to the graphical data. TPQ prepared the graph data and performed the analysis of the resultant RWR scores. HH and TPQ reviewed the literature, designed the experiments, and drafted the manuscript. All authors helped conceptualize the project and revise the manuscript.

